# Further Theoretical Considerations for Next-Generation Proteomics

**DOI:** 10.1101/2021.06.12.446585

**Authors:** Magnus Palmblad

**Author notes:** This article is a follow-up to my recent paper in the *Journal of Proteome Research.

## Abstract

In a recent *Journal of Proteome Research* [paper](https://pubs.acs.org/doi/10.1021/acs.jproteome.1c00136), I described some general properties and constraints of a hypothetical next generation of proteomics technology based on single-molecule peptide sequencing. This work prompted many interesting questions, both from the reviewers of the initial manuscript and later from readers and colleagues. This follow-up paper addresses some of questions by clarifying the original results, considering alternative metrics, and a number of new simulations. Specifically, the discriminative power of individual amino acids is revisited, simulating additional proteolytic agents. These simulations show allowing missed cleavages generally increases the discriminative power of the amino acids in the proteolytic motif. Additional simulations show the effect of non-ideal conditions modelled on the number of proteins lacking proteotypic reads is very small, and that the average number of proteotypic reads per protein follow the same rule on the performance of the optimal choice of labeled amino acids as the number of distinguishable proteins in NeXtProt. The goal of this paper is to expand prior results and continue the scientific discussion on the possibilities of future proteomics technologies.

## Introduction

Among the most rewarding experiences one can have publishing in science is when challenged by a reviewer with an unexpected question from a different perspective. Such questions makes researchers see their work in a different light, often suggesting new directions of exploration. This is exactly what happened in the process leading up to a recent publication in the *Journal of Proteome Research*.^1^ One of the (anonymous) reviewers challenged the generality of the author’s interpretation of the results with what turned out to be an interesting counterexample. After the paper was published, others have asked similar questions on the same analysis of the discriminative power of different amino acids in single-molecule peptide sequencing. This follow-up is an attempt to address these questions and provide a more general interpretation of the results in the published paper.

In the paper, the *discriminative power* of the amino acids was defined by their coefficients in a linear model predicting the total number of discriminated proteins (the number of proteins in NeXtProt with at least one proteotypic peptide-partial read match, or PPRM) as a function of which amino acids are labeled or readable. The coefficients are computed by minimizing the sum of the squares of the differences between the calculated number of discriminated proteins and that predicted by the linear function

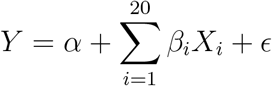

where *Y* is the number of discriminated proteins, *X*_*i*_ = (*x*_*i*1_, *x*_*i*2_, …, *x*_*ip*_)^T^ the vector encoding in which of the *p* simulations amino acid *i* is labeled (*x*_*ij*_ = 1 when amino acid *i* is labeled in simulation *j* and *x*_*ij*_ = 0 when it is not), *α* the intercept, *β*_*i*_ the amino acid coefficients and *ϵ* the residuals. This is known as linear least squares regression.

Similar linear models have been used to predict peptide chromatographic retention based on amino acid composition. ^2,3^ However, the binary nature of the data – an amino acid being either labeled or not – makes the model matrix rank deficient when considering only one number of labeled amino acids, as any column *X*_*i*_ can be written as

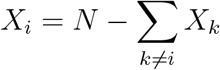

where *N* is the number of labeled amino acids. In other words, we have only 19 and not 20 linearly independent variables in the data. In the paper, this was overcome by varying the number of labeled amino acids and looking at the average discriminative power with 4, 5 or 6 labeled amino acids. Other ways of calculating discriminative power, such as the average number of proteins with proteotypic PPRMs when an amino acid is labeled, would be possible to compute also for a fixed number of labeled amino acids.

The main point of Figure 2D in the original paper was to show that the value of labeling a particular amino acid scales with the frequency of that amino acid in the proteome, i.e. the more frequent the amino acid, the more valuable it is for discriminating between proteins. The linear regressions typically have a large intercept (19,421 proteins for the data shown in Figure 2D in the original paper). Individual coefficients can be negative, albeit small. To facilitate interpretation, the coefficients of the individual amino acids (*β*_*i*_) were plotted including the intercept (*α*) against amino acid frequency. *α* + *β*_*i*_ is the number of predicted discriminated proteins when only amino acid *i* is labeled. These are not necessarily accurate predictions, as the model was trained on data with four to six concurrently labeled amino acids. But they provide meaning to the numbers and a suitable scale for comparisons. Although there are other ways to define the power of (protein) discrimination of the individual amino acids when revealed by a sequencing experiment, they will almost certainly correlate positively with amino acid frequency.

The originally submitted manuscript argued the relatively low values for arginine and lysine by their definition of the cleavage specificity of trypsin. Even with one missed cleavage allowed, the contribution of these residues to peptide and protein discrimination is constrained. We also already have the information that the peptide C-termini is either arginine or lysine (except for the peptides reaching the protein C-terminus). Pinning down which was recognized by trypsin adds less information than labeling another amino acid of similar frequency in the proteome. One of the (anonymous) reviewers asked whether this is generally true, e.g. would it also hold for methionine in CNBr hydrolysis? After some quick simulations, it became clear this is not the case, and adjusted my interpretation accordingly, stating only that the lower discriminative power of arginine and lysine relative to similarly frequent amino acids “may in part be due to these residues defining the cleavage site, and as only one missed cleavage site was allowed”. To better understand how the cleavage motif influences the discriminative power of the amino acids, this follow-up includes additional results from simulating trypsin, CNBr and two additional proteases, allowing for zero or one missed cleavage, using the same code as in the previous paper.

Another recurring question concerns Figure 3 in the published paper, specifically what the general picture looks like, as the figure only compared simulations under ideal and non-ideal conditions for the first 20 simulations. This follow-up therefore shows the full set of matching ideal and non-ideal simulations, indicating how similar they are under the metric of the number of proteins in NeXtProt lacking unique PPRMs, and discuss the limitations of these analyses.

A third question relates to rule #1, stating that the best performance of *n* is similar to the average performance of *n* + 1 labeled amino acids, and how general this rule is. Does it apply to other metrics? How accurate is it? Here we will look deeper at the simulation results with respect to other possible experimental design targets, including the number of unique peptide reads and the average number of proteotypic PPRMs per protein.

Lastly, a concern was raised that RNA-Seq only works because the dynamic range of transcripts is 10^2^-10^3^, implying the comparison between next-generation proteomics and RNA-Seq would be flawed, as most proteomes span more orders of magnitude in protein abundance. Though the comparison is imperfect, the premise for this particular argument is false, as shown in RNA-Seq studies. ^4–7^ Even with a conservative minimum number of reads, the dynamic range of such RNA-Seq experiments can be 10^5^ in technology that is in routine use today. Older technologies such as microarrays^8^ have a smaller dynamic range - on the order of 10^3^. For an accessible review of the fundamentals and history of transcriptomics methods, see Lowe et al.^9^ Given how rapidly in particular next-generation sequencing has evolved in recent years, there is little to suggest these technologies would have already reached their maximum potential. In their review of proteomics beyond mass spectrometry, Timp and Timp^10^ discuss dynamic range challenges and possible solutions in current and future proteomics technology in more detail, as do Alfaro et al. in their recently published and insightful perspectives. ^11^ Whether or not we wish to store all the redundant raw data corresponding to the same peptide sequence, or implement triggers for storing some but not all redundant sequences, or only entirely distinct sequences, is a separate engineering problem with a precedent in particle physics.^12^

First, however, we will set the stage by returning to the terminology introduced in the paper and provide some additional explanation for the terms and acronyms.

## About the nomenclature

The published paper focused on matching partial (tryptic) peptide reads to proteins in NeXtProt, where partial refers to the possibility that only some of the 20 proteinogenic amino acids are *readable*, i.e. directly revealed in the sequencing experiment. The term peptide-partial read match, PPRM, was introduced as a next-generation proteomics analog of peptide-spectrum matches. A shorter “PRM” for peptide-read match could also have been chosen. However, this acronym is already in frequent use in proteomics for parallel reaction monitoring.^13^ One possible drawback of the “partial” in PPRM is that it makes the term less fitting when all 20 amino acids are readable, though it may still be the case that individual reads are truncated and therefore still partial, albeit in a different sense. Such truncated reads would of course have to be dealt with in the analysis of the data. In summary therefore, peptide-partial read match appears to be a term generally applicable in next-generation proteomics as discussed. In protein identification, *proteotypic* peptide (reads) are particularly valuable, whether generated by mass spectrometry or any other technology. The trivial acronym for proteotypic partial peptide read matches would be PPPRM. This could be further abbreviated to P3RM, analogous with the Mitogen Activated Protein (MAP) kinase kinase kinase, MAP3K.

In the analysis of the simulation results and as further explained above, the previous paper also looked at the *discriminative power* of the amino acids. This usage agrees with its generic connotation of how effective a particular discriminator (here the pattern based on reading a particular amino acid) is in its ability to correctly categorize items (the peptides being the items and the proteins the categories).

## Results

### Amino acid discriminative power

As it turns out, the original manuscript statement that the amino acids found in the proteolytic motif are less informative than other amino acids of similar frequency in the proteome, is not universally true, including when missed cleavages are allowed (Figure 1 below). A clear counterexample is methionine in CNBr, which is at least as informative as other residues of similar frequency when a single missed cleavage is allowed (Figure 1B below). However, when no missed cleavage is allowed, the statement is true for the investigated proteolytic agents trypsin, CNBr, neutrophil elastase and Asp-N regardless the frequency of the amino acids, whether cleavage is C- or N-terminal, or the number of amino acids in the motif (Figure 1). For the proteases, the amino acid in the motif are below the average for amino acids of similar frequency. But the effect is far smaller than when no missed cleavages are allowed. The regression for trypsin with one missed cleavage allowed in Figure 1a below is similar to Figure 2D in the original paper, despite being based on different random amino acid selections. The general rule (rule #3 in the original paper) that the discriminatory power of the amino acids scales with their frequency in the proteome holds for all simulated proteolytic agents.

**Figure 1:**
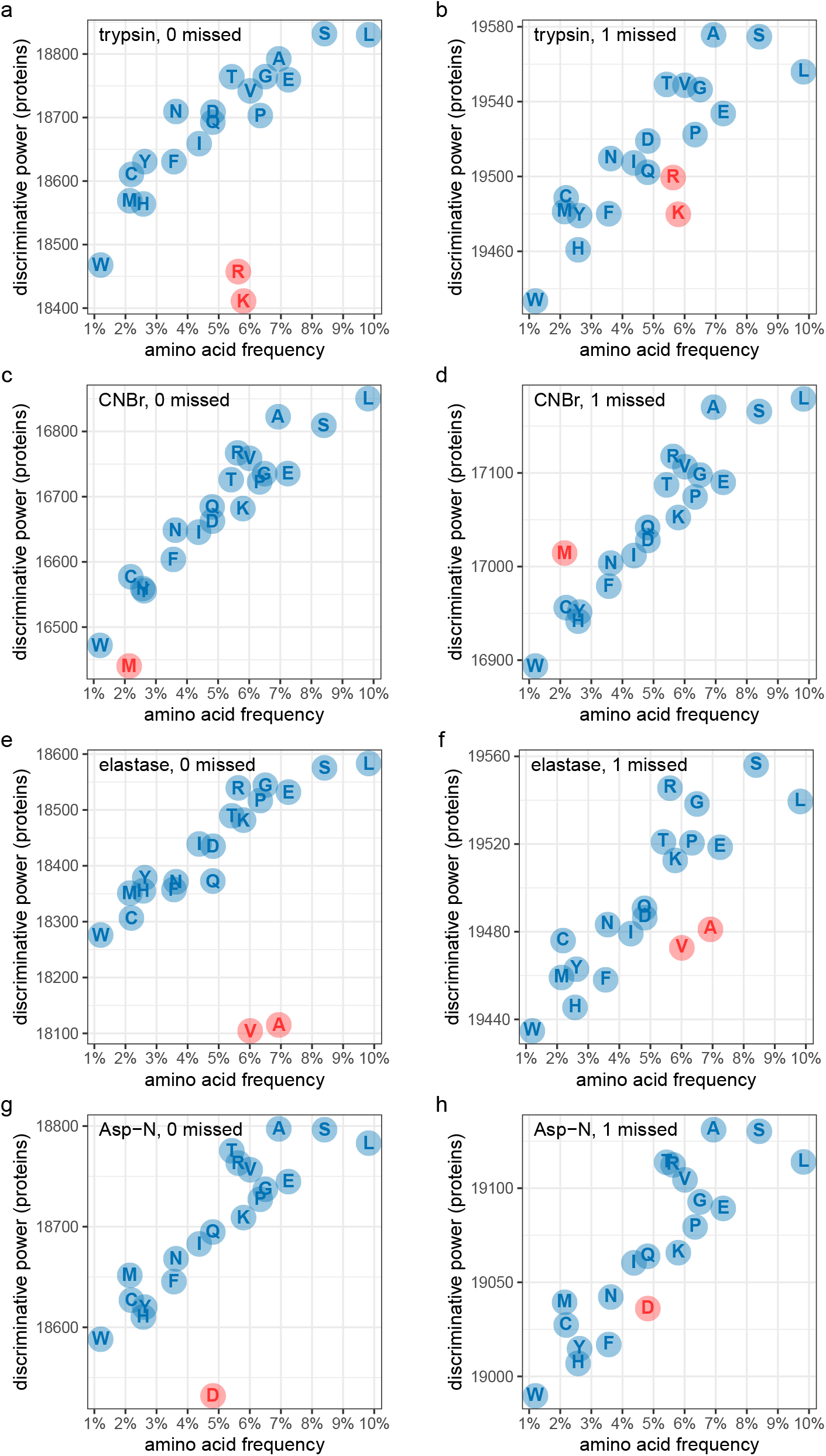
Power of the amino acids to discriminate NeXtProt sequences by tryptic (a,b), CNBr (c,d), neutrophil elastase (e,f) and Asp-N (g,h) peptides, with zero or one missed cleavage allowed. Amino acids in the proteolytic motif are shown in red.

### Ideal *vs* non-ideal conditions

Figure 2 shows the results comparing the full matching sets of 400 simulated random selections of 1 to 20 amino acids for the ideal and non-ideal conditions. Obviously, the selection of 20 out of 20 amino acids will always be the same, and there are no gaps of uncertain length in the reads. There is very little difference in the number of proteins lacking proteotypic PPRMs caused by gap length uncertainty in the non-ideal case. However, the number of possible peptide reads increases dramatically (right panel). This simple modelling of non-ideal conditions most informative when few amino acids are labeled and the gaps dominate the reads.

**Figure 2:**
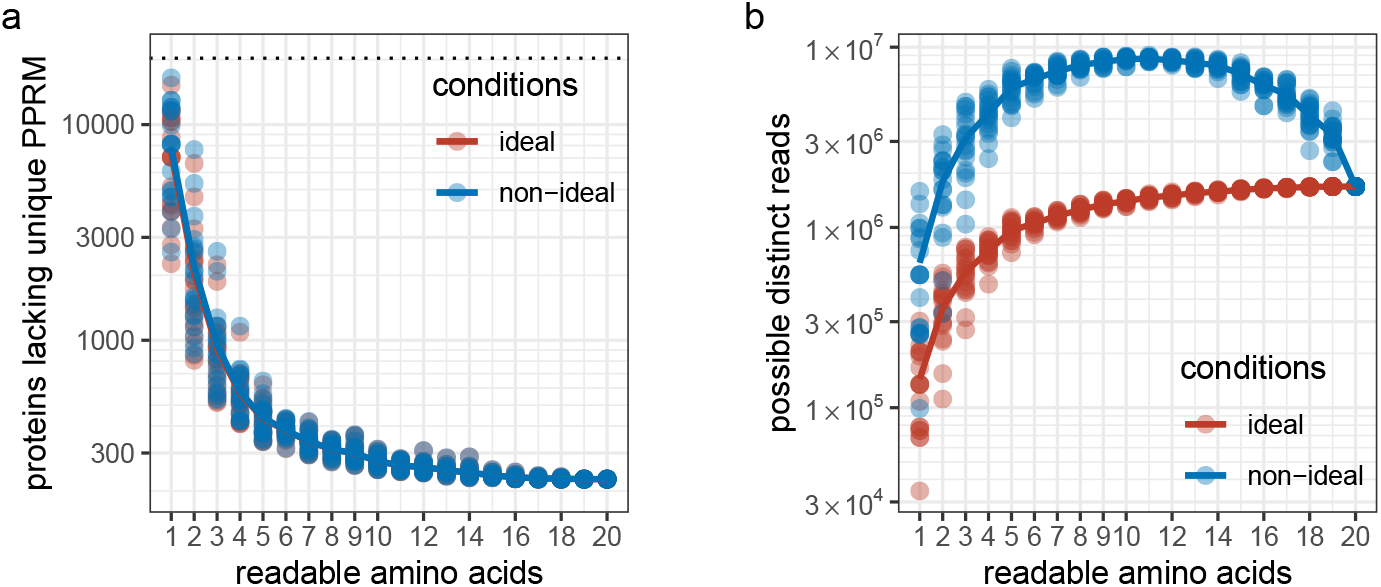
Simulation (a) of matching tryptic partial peptide reads to NeXtProt proteins under ideal (red) and non-ideal (blue) conditions, for a maximum peptide length of 50, completing the data in Figure 3 in the original paper. Under this simplistic modelling, the number of proteins lacking unique PPRMs is not dramatically changed, at least with three or more amino acids made readable. As already stated in the original paper, the number of possible reads (of any length) is around six-fold higher when labelling 4 to 10 amino acids (b). When more amino acids are readable, the simplistic model of non-ideal conditions is less relevant, as there are fewer gaps in the reads contributing to ambiguity in the model.

### Generality of rule #1

The first of the three rules in the original paper summarized observations from the simulations as the best selection of *n* amino acids performing very similar to the average selection of *n*+1 amino acids. How general is this observation? Is it limited to the a particular metric, such as the number of proteins with at least one proteotypic PPRM, or does it hold for other metrics as well? How similar is “similar”? Figure 3 below shows the results from simulating 400 random selection of 1 to 20 amino acids labeled with maximum tryptic peptide read with maximum peptide lengths of 10, 20 and 50 amino acids, allowing one missed cleavage. We see that the rule also holds approximately for these metrics.

How accurate is the rule? Trivially, it holds exactly for *n* = 19, as there is no difference in information content (under ideal conditions) between labeling 19 or 20 amino acids, as the unlabeled amino acid fills all gaps in the read. In the more relevant range *n* = 4, 5 or 6, the deviation between the best performing selections of *n* and average selections of *n* + 1 amino acids is about 10%. In the simulations shown in the original paper the differences were 9.9% (*n* = 4), 6.4% (*n* = 5) and 11.6% (*n* = 6) for read lengths up to 10 amino acids, 0.5 % (*n* = 4), 1.5% (*n* = 5) and 7.0% (*n* = 6) for reads up to 20 amino acids and 5.4% (*n* = 4), 11.2% (*n* = 5) and 7.4% (*n* = 6) for reads up to 50 amino acids. These numbers could be compared with the differences between the best and worst performing selections of *n* amino acids, which is typically twofold or more (for *n* = 4, 5 or 6). While the averages are stable, it is possible some extreme values will change slightly with additional simulations. However, the rule is sufficiently precise for a simple heuristic in the design of single-molecule peptide sequencing methodology: if given free choice in which amino acids to label, we can expect performances similar to those with one more amino acid labeled but when the choice is constrained.

**Figure 3:**
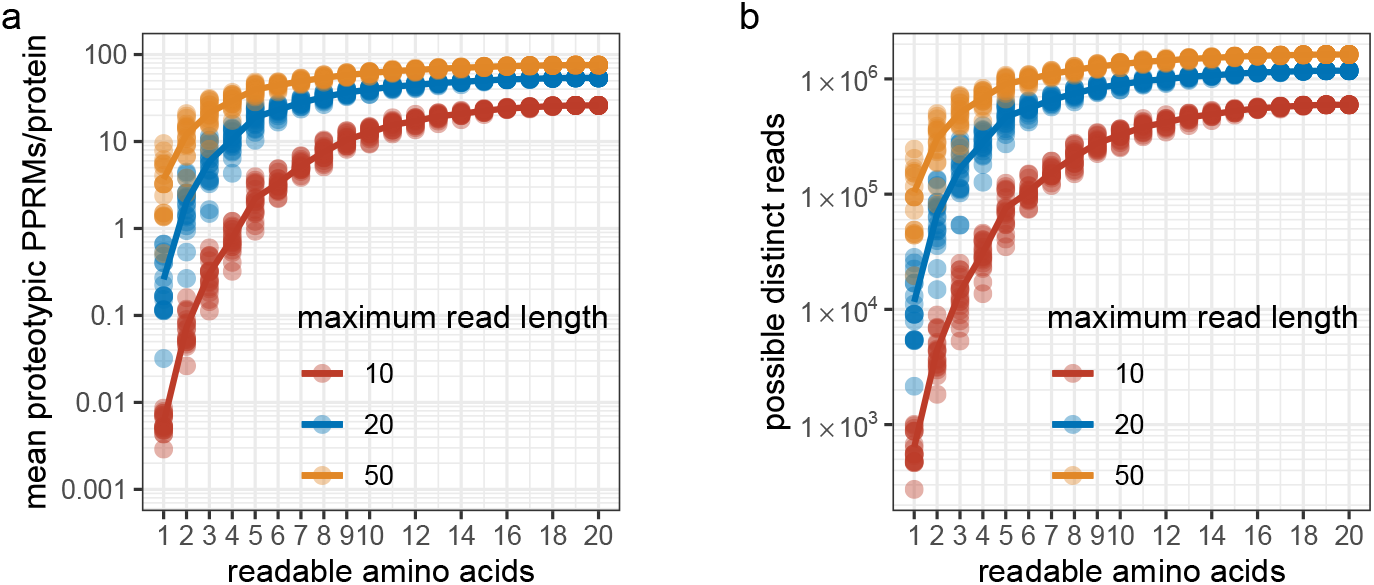
The average number of proteotypic PPRMs per protein in NeXtProt (a) and the total number of possible unique end-to-end reads of peptides with a maximum of 10, 20 and 50 amino acids (b) as function of the number of labeled amino acids (*cf* Figure 2A in the original paper). To place these numbers in some context we can note that for the most frequent amino acids (Leu, Ser, Glu), around half of the 2^1^ + 2^2^ + … + 2^10^ = 2^11^ 1 1 = 2046 possible sequences of lengths 1 to 10 are found in NeXtProt. With two amino acids labeled, at most around 11,000 of the 3^1^ + 3^2^ + … + 3^10^ = 88,572 possible sequences can be generated by end-to-end tryptic peptide reads from NeXtProt. Note that these simulations ignore shorter, truncated, reads of longer peptides, i.e. we assume we only analyze end-to-end reads, with each end either corresponding to a proteolytic cleavage site or protein terminus. This assumption is somewhat analogous to knowing the mass on the intact peptide in mass-spectrometry based proteomics, i.e. we know when we have read to the end of the peptide sequence and assume this end corresponds to either a cleavage site or protein terminus. It is possible, perhaps even likely, that in single-molecule protein sequencing, we will generate and wish to use truncated reads, with neither or only one of the ends corresponding to a cleavage site or protein terminus.

### Conclusions

Each single-molecule sequencing technology will likely have its own advantages and short-comings. The models used here are simple and intended to convey general messages rather than optimize the design of specific experiments. The modelling of non-ideal effects is only a first attempt toward understanding how sensitive peptide-partial read matching would be to read errors, such as the modelled gap extensions. Each single-molecule sequencing technology will have its own particular confusion matrix that, when known, could be used in false-discovery rate estimation for PPRMs and proteins from single-molecule sequencing data. Another conclusion that can be drawn from the extended analysis of amino acid discriminative power is that if the proteolysis is driven to completion, there is little value in labeling the amino acids occurring in the proteolytic motif, regardless of agent.

In deciding to upload these comments, explanations and additional analyses to bioRxiv, the intention and hope is that they will serve as a starting point for a conversation among those interested in this emerging topic. Certainly I look forward to learn from and be inspired by these discussions on future proteomics technologies.

## Experimental

### Simulations

The simulations here were based on the same R code as the original paper, changing the `enzym` and `missedCleavages` arguments in the `cleave`() function, and the metrics captured. To compare different proteolytic agents, the same set of 100 random selections of each of 4, 5 and 6 amino acids where used for each of the proteolytic agents trypsin (with the Keil rules^14^), CNBr, neutrophil elastase and Asp-N endopeptidase, and each number of allowed missed cleavage sites (zero or one). The selections were different than those used for trypsin in the original paper. The non-ideal conditions were simulated for the same 400 random selections of 1 to 20 amino acids as used in the simulations of ideal conditions in the original paper, with the same limits in peptide length (50) and number of missed cleavages allowed (one).

### Analysis

The linear regression of the number of discriminated proteins as a function of labeled amino acids were performed with `lm()`as in the original work, specifically comparing the different proteolytic agents with zero and one missed cleavage allowed. The previous simulation results from trypsin under ideal conditions, with 1 missed cleavage allowed and a maximum peptide length of 50 amino acids were compared with respect to the total number of unique peptide reads as well as the average number of proteotypic PPRMs per protein, both as functions of the number of labeled amino acids.

## Code availability

Both the simulation and analysis code, as well as all simulation results, are available on GitHub under the Apache License 2.0: https://github.com/magnuspalmblad/NGP.

## Acknowledgement

The author thanks the anonymous reviewers of the original paper, Mitra Ebrahimpoor for input on linear regression in R, Jeroen Laros and Leon Mei for discussions on RNA-Seq, and finally Benjamin Neely and Rob Marissen for their helpful comments on the manuscript.

